# A quantitative genetic model for indirect genetic effects and genomic imprinting under random and assortative mating

**DOI:** 10.1101/2024.05.08.593214

**Authors:** Ilse Krätschmer, Matthew R. Robinson

**Affiliations:** Institute of Science and Technology Austria, Klosterneuburg, Austria

**Keywords:** quantitative genetics model, epigenetic, parent-of-origin, indirect effects

## Abstract

An individual’s phenotype reflects a complex interplay of the direct effects of their DNA, epigenetic modifications of their DNA induced by their parents, and indirect effects of their parents’ DNA. Here, we derive how the genetic variance within a population is changed under the influence of indirect maternal, paternal and parent-of-origin effects under random mating. We also consider indirect effects of a sibling, in particular how the genetic variance is altered when looking at the phenotypic difference between two siblings. The calculations are then extended to include assortative mating (AM), which alters the variance by inducing increased homozygosity and correlations within and across loci. AM likely leads to covariance of parental genetic effects, a measure of the similarity of parents in the indirect effects they have on their children. We propose that this assortment for parental characteristics, where biological parents create similar environments for their children, can create shared parental effects across traits and the appearance of cross-trait AM. Interestingly, the genetic variances is increased under AM for the childmother-father design, while it is decreased for the sibling difference. Our results demonstrate that it is not possible to get unbiased estimates of direct genetic effects without controlling for parental and parent-of-origin effects.

## Introduction

In humans, parental influence persists long beyond childhood, shaping adult health and disease^1^. A basic principle of genetics is that observable characteristics result from two sources: the expression of an individual’s DNA and the environment that they experience^2^. Parental characteristics shape a child’s environment, giving rise to indirect genetic effects, where the genotypes of both the mother and father influence the traits of their children, beyond simply the alleles that are inherited^2^, as displayed in Figure 1. Many studies have now shown that failing to separate the effects of an individual’s DNA from the indirect effects of their parental genotypes results in confounding within genetic association studies^3,4,5,6^. Moreover, it has been shown that maternal effects cause the appearance of parent-of-origin effects that can mimic genomic imprinting, and vice versa^7^.

**Figure 1:**
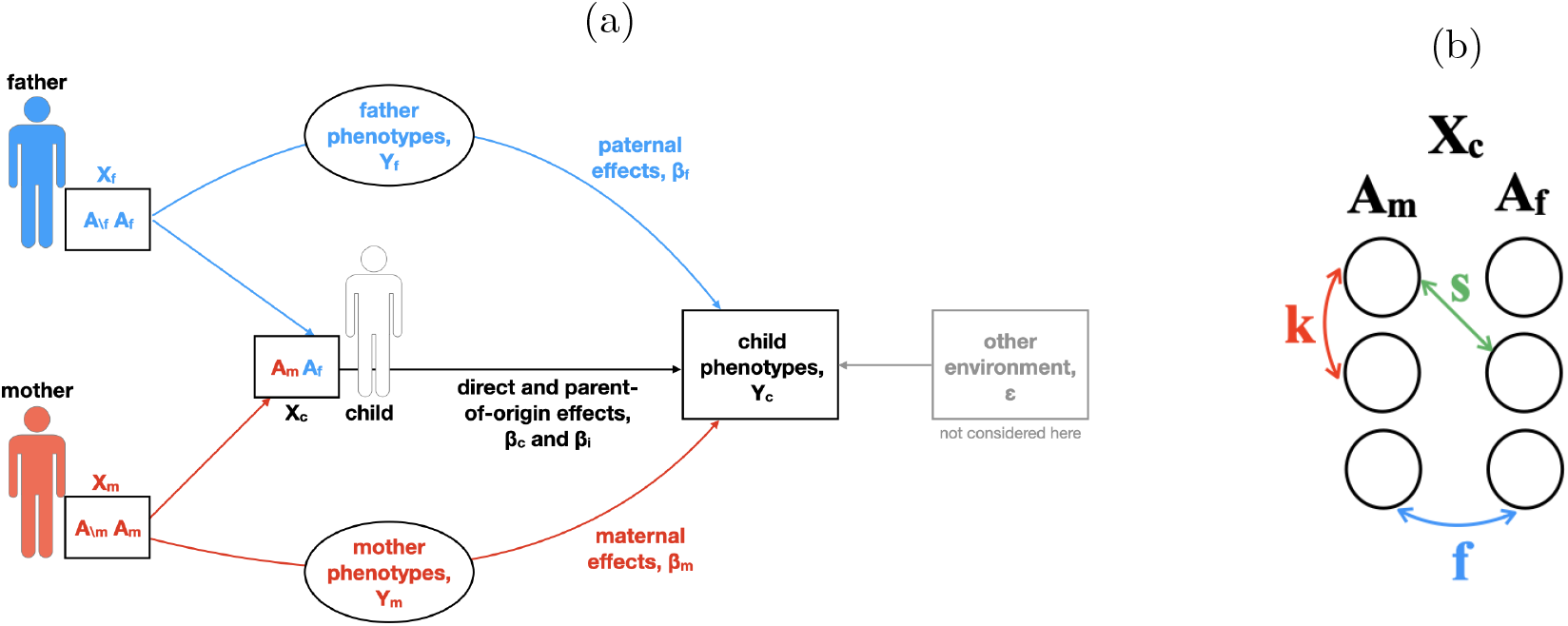
Definition of variables. (a) Direct (***β***_***c***_), parent-of-origin (***β***_***i***_) and indirect maternal (***β***_***m***_) and paternal (***β***_***f***_) genetic effects on the phenotypes of children (***Y***_***c***_). The parental genotypes (*X*_*m*_, *X*_*f*_) are split into transmitted alleles (*A*_*m*_, *A*_*f*_), which constitute the genotype of the child (*X*_*c*_), and untransmitted ones (*A*_*\m*_, *A*_*\f*_). Parental effects can be mediated through any parental phenotype (*Y*_*m*_, *Y*_*f*_). The variable ***ϵ*** Represents variations created by other environmental effects experienced by the children. (b) Correlations within and between loci introduced through AM. The child’s DNA (*X*_*c*_) consists of maternally inherited alleles (*A*_*m*_) and paternally inherited ones (*A*_*f*_). The correlation between alleles at a locus is given by *f*, the one between alleles at different loci within the same gamete by *k* and the one between pairs of *A*_*m*_ and *A*_*f*_ alleles at different loci by *s*.

Previous quantitative genetics models derived the genetic population variance under the influence of maternal effects^8,6^, or imprinting^9^ or both^10,7,11^ under random mating. *Wolf et*.*al*. demonstrated how indirect maternal effects contribute to the variance under inbreeding^8^. Other studies^12,13,14^ provided a general framework to show the influence of confounding in family-based genome-wide association studies (GWAS), and how polygenic risk scores (PGS) are biased under assortative mating in the presence of indirect parental effects.

But so far, none of the models considered direct, indirect maternal and paternal, and parent-of-origin genetic effects jointly. Here, we first derive the genetic variance for a single locus under random mating assuming that all four effects (direct, indirect maternal and paternal, and parent-of-origin) influence an individual’s phenotype. We then extend the model to include also the indirect effects of a sibling and consider another common family-based design in human studies: the phenotypic difference between two siblings. Next, we show how the genetic variance changes in the presence of assortative mating (or inbreeding). Lastly, we demonstrate how to include multiple loci in the model. We then also verify our theoretical model with forward-in-time simulations, based on genotypes from the 1000 Genomes project.

Though we primarily consider humans and use the corresponding terms, we want to point out that our calculations are not limited to humans, but generally applicable to diploid sexually reproducing species that rear their offspring.

### Single-locus model under random mating

We first consider a simple single autosomal locus with two alleles (*A*_1_ and *A*_2_) with frequencies *q*_1_ and *q*_2_ that can have the following four effects on the child’s phenotype: direct (*β*_*c*_), indirect maternal (*β*_*m*_), indirect paternal (*β*_*f*_)and parent-of-origin (imprinting, *β*_*i*_). The alleles of the child at the locus are ordered, referring first to the maternally inherited allele and second to the paternally inherited one, i.e. *A*_*m*_*A*_*f*_. The genotypes *X*_*c*_, *X*_*m*_, *X*_*f*_, are coded as 0, 1 and 2 for *A*_1_*A*_1_ homozygotes, *A*_1_*A*_2_ or *A*_2_*A*_1_ heterozygotes and *A*_2_*A*_2_ homozygotes, respectively. We use subscripts m, f, c, and i to denote maternal, paternal, direct (child) and parent-of-origin (imprinting) effects, genotypes and genotypic values. For heterozygous children, the parent-of-origin effect is positive for maternally inherited *A*_2_ and negative for paternally inherited *A*_2_. From these definitions, the possible genotypes, *X*_*c*_, and phenotypes, *Y*_*c*_, of the child given the genotype of the parents and imprinting, are shown in Table 1. The phenotypic values represent the deviations from the case where both parents and child have *A*_1_*A*_1_ genotypes.

**Table 1:**
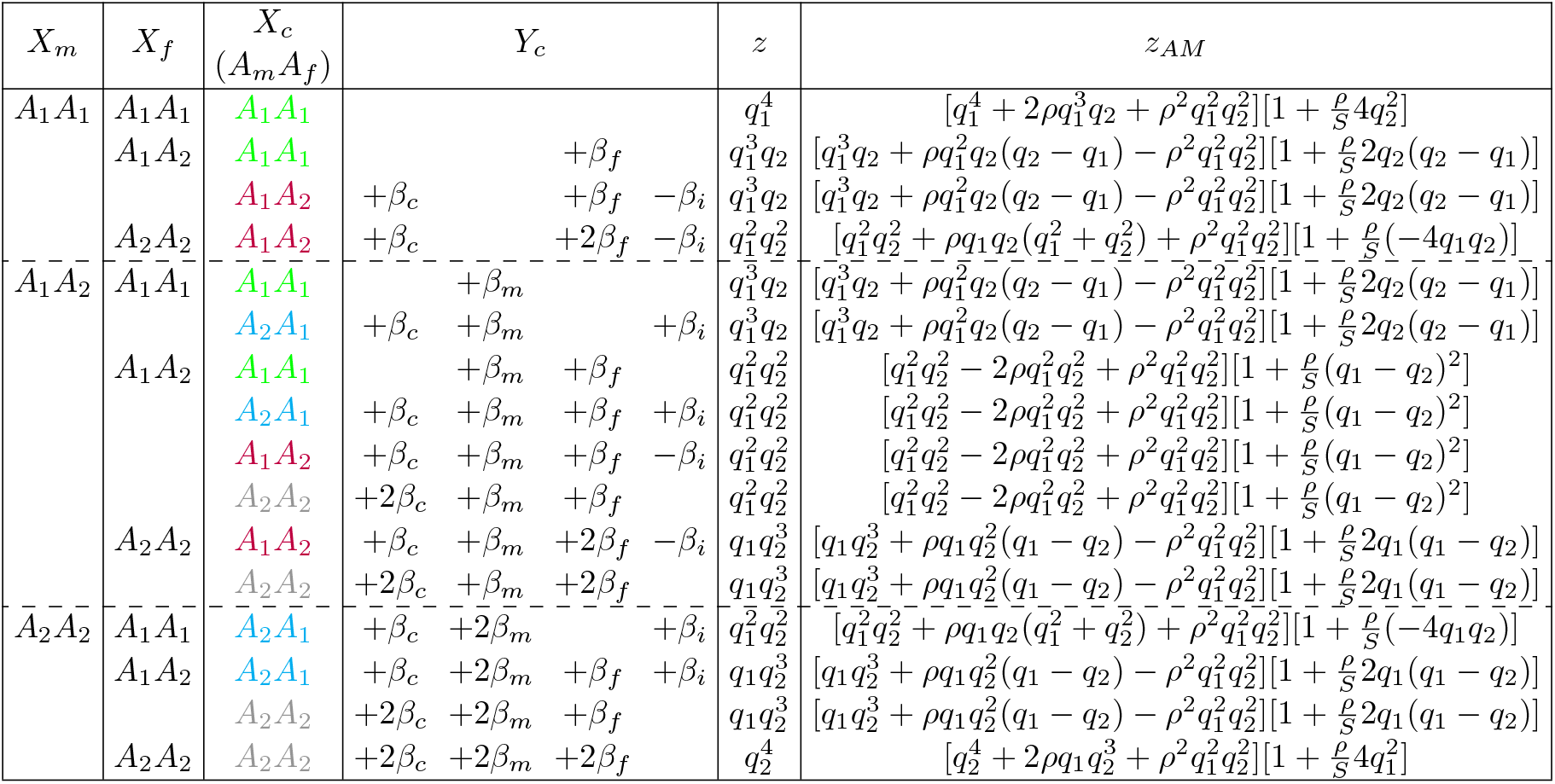
A simple single-locus model with indirect parental and parent-of-origin genetic effects. Maternal (*X*_*m*_), paternal (*X*_*f*_) and child’s genotypes (*X*_*c*_), phenotypes of the child (*Y*_*c*_), mating frequencies under random mating (*z*) and under assortative mating (*z*_*AM*_) taken from Ref.^15^ where *S* = 2*q*_1_*q*_2_(1 + *ρ*). Parental heterozygous genotypes are not ordered and are used interchangeably. Refer to text for more details.

Table 1 shows that indirect parental genetic effect genotypic values, *X*_*m*_*β*_*m*_ and *X*_*f*_ *β*_*f*_, and parent-of-origin effect genotypic values, *X*_*i*_*β*_*i*_, are confounded due to the fact that not every child genotype can be produced by all parental genotypes. For example, parents who are both the same type of homozygote (both either *A*_1_*A*_1_ or *A*_2_*A*_2_) will only have *A*_1_*A*_1_ and *A*_2_*A*_2_ children. Each of the two types of homozygous parents (*A*_1_*A*_1_ and *A*_2_*A*_2_) can produce only one of the two types of reciprocal heterozygote (*A*_1_*A*_2_ and *A*_2_*A*_1_). The children of *A*_1_*A*_1_ fathers all have +*β*_*i*_, with the exception of *A*_1_*A*_1_ children whose parents are both *A*_1_*A*_1_. The children of *A*_2_*A*_2_ fathers all have *−β*_*i*_, with the exception of *A*_2_*A*_2_ children whose parents are both *A*_2_*A*_2_. The reverse is true for mothers, where the children of *A*_1_*A*_1_ mothers all have *−β*_*i*_, with the exception of *A*_1_*A*_1_ children whose parents are both *A*_1_*A*_1_, and the children of *A*_2_*A*_2_ mothers all have +*β*_*i*_, with the exception of *A*_2_*A*_2_ children whose parents are both *A*_2_*A*_2_. This clearly shows that when there are indirect parental genetic effects, the phenotypes of the reciprocal heterozygotes will differ, creating the appearance of parent-of-origin effects. Likewise, differences between reciprocal heterozygotes due to parent-of-origin effects will appear as parental genetic effects.

Table 1 also provides the expected probability of the different parent-child combinations within the population (mating frequency), *z*, under random mating and Hardy-Weinberg equilibrium, where (*q*_1_ + *q*_2_) = 1 = (*q*_1_ + *q*_2_)^2^, as well as the probabilities under assortative mating, *z*_*AM*_, which will be discussed in the next section. The phenotypic values, *Y*_*c*_, and expected probabilities, *z*, in Table 1 can be used to calculate the mean and variance contributed by direct, indirect parental genetic and parent-of-origin effects within the human population under random mating. The population mean is given by

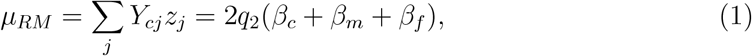

where *j* loops over all possible genotypes of the child. The mean is not influenced by the parent-of-origin effects, as they are on average zero. The genetic variance for a single locus is:

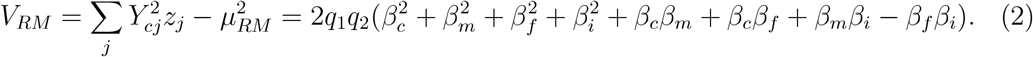

More details on the calculations are provided in the Supporting Information.

Equation 2 shows that direct genetic effect genotypic values covary with both maternal and paternal genetic effect genotypic values, which in turn covary with parent-of-origin effect genotypic values, creating a greater degree of potential confounding between all four components than has been previously considered. The additive genetic variance among child’s genotypes, apart from the expected terms due to direct effects 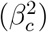, also contains components contributed by parental genetic effects 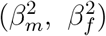, direct-parental genetic covariance (*β*_*c*_*β*_*m*_,*β*_*c*_*β*_*f*_), parent-of-origin effects 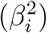, and parental-parent-of-origin genetic covariance (*β*_*i*_*β*_*m*_, *β*_*i*_*β*_*f*_). Parent-of-origin effects contribute not only directly, but also dependent on their covariance with parental genetic effects, i.e.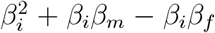, which represents the degree to which loci with parental genetic effects are also imprinted loci in the child. Note the negative sign of *β*_*i*_*β*_*f*_, which is due to the definition of parentof-origin effects (positive for the maternally inherited allele and negative for the paternal one).

The implications of Equation 2 are that phenotypic variation can be attributable to the single nucleotide polymorphism (SNP) markers of an individual (i.e., SNP heritability 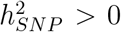) and marker-phenotype associations can be discovered in population studies, but the underlying pattern of causality can be anything from: (i) the locus does not directly affect the expression of the trait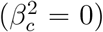, but instead only has an effect in one or both of the parents; to (ii) both direct and parental genetic effects may be present at a locus, but the parental effects may cancel out if they covary negatively; or (iii) any combination of terms and their covariance.

### Differences in siblings

The model presented in the previous section can be extended to include genetic effects of a sibling on the child’s phenotype, additionally to direct, indirect parental and parent-of-origin genetic effects. The derivations of population mean and variance including sibling effects (*β*_*s*_) are given in the Supporting Information. However, it is more interesting to look at the relationship of the phenotypic and genotypic differences in siblings as the difference in siblings provides an estimate of direct genetics effects unbiased of indirect parental effects^16,17,18^. Typically, sibling analyses do not consider parent-of-origin effects, nor the indirect genetic effects from siblings. Using the phenotypic differences between child and sibling, (*Y*_*c*_*−Y*_*s*_), and the expected probabilities of the genotypes for the different sibling combinations, which are given in Table 2, we can calculate the expected mean and variance at population level. The total mean is 0, while the variance is

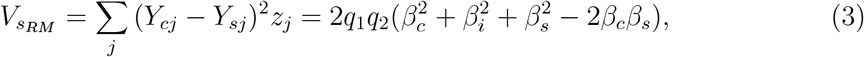

which shows that components due to indirect parental effects do indeed cancel in the difference, but those due to indirect sibling and imprinting effects remain.

**Table 2:**
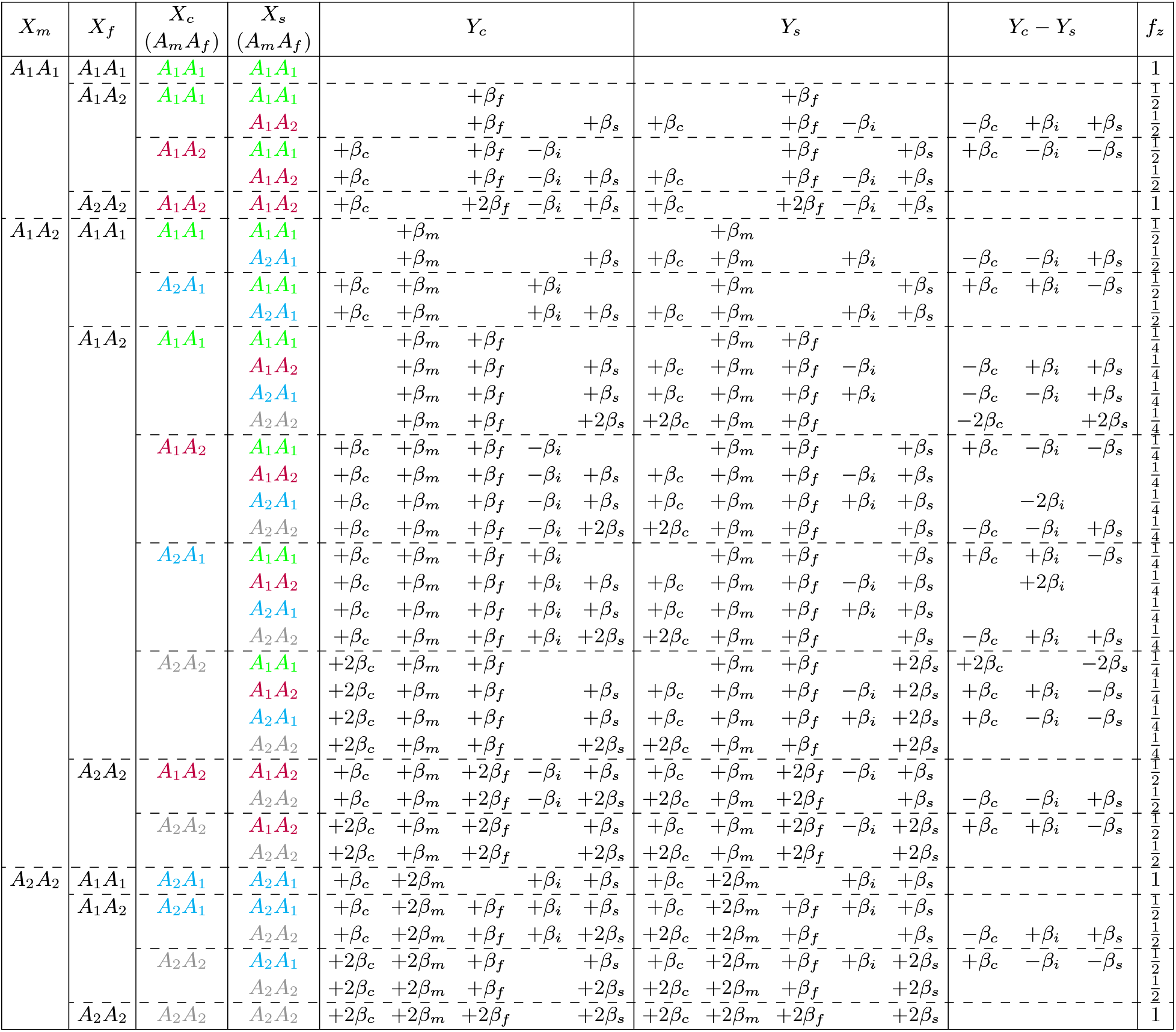
Extension of Table 1 to include siblings and their differences. Mating frequencies are given in Table 1 which have to be multiplied with the factor *f*_*z*_ to account for the different sibling combinations. Parental heterozygous genotypes are not ordered and are used interchangeably.

Importantly, if there is an indirect effect from the sibling on an individual’s phenotype, the variance increases with the squared sibling’s effects, but decreases with the covariance between the individual and its sibling. This results in a confounding of direct and indirect sibling effect genotypic values, which always oppose each other in sign, as shown in Table 2. Thus, any indirect sibling genetic effect will act to alter the variance due to the direct genetic effects. But it does appear possible to separate parent-of-origin effects within this experimental design under random mating.

### Single-locus model under assortative mating

Assortative mating (AM) acts to alter the mating frequencies in the population as it increases parental similarity at trait-associated loci, which is reflected in the correlation, *ρ*, among the parents in their alleles. As a consequence of AM, the genotype frequencies are changed, and thus also the probabilities of parents producing offspring of a certain genotype, given as *z*_*AM*_ in Table 1. But the minor allele frequencies in the population remain invariant. The altered offspring frequencies, *z*_*AM*_, are taken from Ref.^15^. This modification leaves the population mean unchanged under AM, but it alters the variance.

Under assortative mating, we find that a single locus with direct, indirect parental and parent-of-origin effects makes the following contribution to the genetic variance:

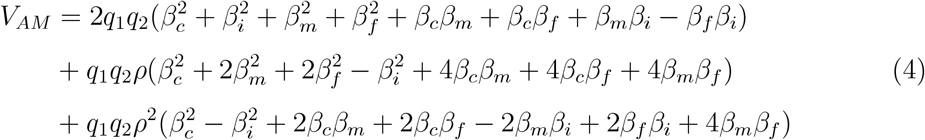

The *ρ* term reflects a correlation among mates at causal SNP variants (or those correlated with an underlying causal variant) and is equivalent to the inbreeding coefficient, *F*. Equation 4 shows that the increase in the variance contributed by a locus under AM is dependent upon the relationships among the direct, parental genetic and parent-of-origin effects.

In contrast, we find that the single-locus genetic variance under assortative mating for the sibling difference is reduced by the parental correlation, *ρ*:

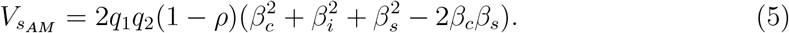

More details can be found in the Supporting Information.

### Multi-locus model

We now extend our model from a single locus to multiple loci (assuming a polygenic additive genetic model) by summing over all *l* causal variants and taking into account correlations, *k*_*ij*_, between loci. The total genetic variance under random mating is

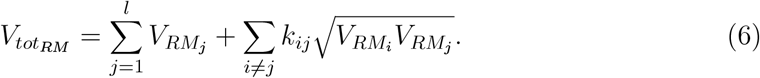

The covariances between loci, given in the second term of Equation 6, are dependent upon the correlations among the causal variants and the concordance in their effects size.

Extending the single-locus model for differences in siblings to multiple loci can be done through Equation 6, substituting *V*_*RM*_ with Equation 3. We can also easily adapt Equation 6 to include AM by substituting *V*_*RM*_ with *V*_*AM*_, or in case of the sibling difference *V*_*sAM*_, where AM-induced changes within a single locus are already taken into account. AM-induced correlations between loci (both cis and trans), on the other hand, will alter the value of the correlation term, *k*_*ij*_.

The change in variance due to AM induced correlations between alleles of mates at a locus and also between loci has been shown by previous theory^19^ and empirical research^20^. An equilibrium is reached under AM, where there is inflated genetic variance, a larger correlation between relatives for traits driving AM and increased homozygosity at causal loci. To put our work in the context of previous theory^19^, we use a similar approach to Ref.^19^ to understand the effect of AM on the variance taking into account multiple loci and parental influences. We follow the approach outlined in Chapter 4.7 of Ref. ^19^, where all alleles of *l* causal loci are assumed to have the same frequencies, *q*_1_ and *q*_2_ = 1 *− q*_1_, and the same effect *α* on the trait under study. In our work, we consider the parental genetic influence as well as direct effects, and thus *α* needs to be replaced, depending upon whether the allele is inherited from the mother or the father. The variance of a single allele from the mother is determined to be *q*_1_*q*_2_(*β*_*c*_ + *β*_*m*_)^2^, assuming that the effect of the allele is either (*β*_*c*_ + *β*_*m*_) or 0, with frequency *q*_1_ and *q*_2_. We temporarily assume that parent-of-origin effects are absent, as the procedure of Ref. ^19^ is based on the variance of single alleles, but we will introduce these effects again later. The mean of the maternal allele is then given by *q*_1_(*β*_*c*_ + *β*_*m*_) + *q*_2_ · 0 = *q*_1_(*β*_*c*_ + *β*_*m*_). The variance is calculated as

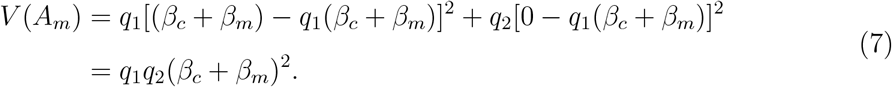

Similarly, the variance for one paternal allele is

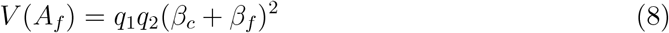

Without any correlations between loci or alleles, the sum of the total variance of an individual, *V* (*X*), is just the sum of the variances over all loci, where one allele at each locus is inherited from the mother and one from the father

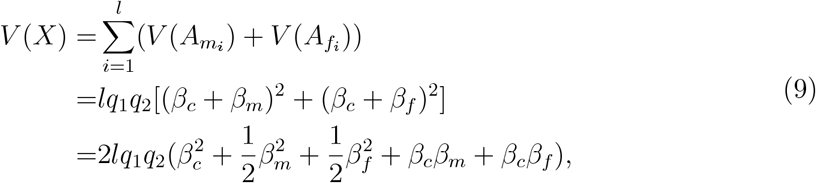

which is the expectation for the additive genetic variance in a randomly mating population in the presence of direct and indirect parental effects.

Through AM, correlations between alleles of mates and between loci are introduced, as shown in Chapter 4.7 of Ref.^19^. Relevant correlations present in the child are correlations between the alleles at the same locus, denoted *f*; at different loci, denoted *s*; as well as at different loci, but within the same gamete, denoted *k*. These are also displayed in Figure 1. Due to the correlations, the total variance then changes to

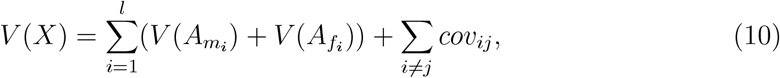

where the covariance term *cov*_*ij*_ consists of the following three parts, adapted from Ref.^19^:

1. The covariance of a pair of *A*_*m*_ and *A*_*f*_ alleles at the same locus for all *l* pairs is given by *lfq*_1_*q*_2_(*β*_*c*_ + *β*_*m*_)(*β*_*c*_ + *β*_*f*_), where *f* represents the correlation between *A*_*m*_ and *A*_*f*_.
2. The covariance between alleles at different loci, but within the same gamete, is *l*(*l −* 1)*/*2 *· kq*_1_*q*_2_(*β*_*c*_ + *β*_*m/f*_), where the correlation is denoted with *k*. The covariance differs for maternal and paternal alleles in the parental effect *β*_*m/f*_.
3. There are *l*(*l −* 1) pairs of *A*_*m*_ and *A*_*f*_ alleles at different loci with correlation *s* and covariance *l*(*l −* 1)*sq*_1_*q*_2_(*β*_*c*_ + *β*_*m*_)(*β*_*c*_ + *β*_*f*_).

These three covariance terms all act to inflate the genetic variance, as is expected from AM. However, AM also induces a change in the mating frequencies, which the first covariance term does not explicitly take into account. Rather, it is assumed that the correlation *f* acts in the same way on all genotypes, which is akin to assuming that the mating frequencies are not changed. Thus, while previous theory has established that an equilibrium is reached under AM, where there is inflated genetic variance, a larger correlation between relatives for traits driving AM and increased homozygosity at causal loci^19^, the increased homozygosity is not directly accounted for when calculating the expected increase in genetic variance.

Our single-locus model under AM presented above incorporates changes in the mating frequency under AM, which accounts for the increase in variability at one locus that is induced by *f* (which we denote as *ρ* above). To extend our model to multiple loci, we therefore only need to consider the covariances across loci. Instead of studying the correlations between a pair of alleles, we look at the correlation *k* between a pair of genotypes at different loci, i.e. (*A*_*m*_*A*_*f*_)_*i*_ and (*A*_*m*_*A*_*f*_)_*j*_. This also enables us to include imprinting effects.

For *k* = *corr*((*A*_*m*_*A*_*f*_)_*i*_, (*A*_*m*_*A*_*f*_)_*j*_), the covariance for *l*(*l −* 1) possible pairs is given by 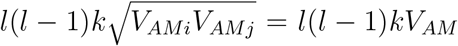, using the variance under AM defined in either Equation 4, or Equation 5 for differences in siblings. Correlations induced by AM between loci will thus inflate the variance by the factor *l*(*l −* 1)*k*. Extending this to the case where the correlations between the loci are not equal and effects and allele frequencies are not the same for each locus, the covariance changes to 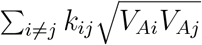. Depending on the sign of the covariance term, it will either increase or reduce the genetic variance, and is thus important to take into account.

In human studies, another common family design is using transmitted and untransmitted parental alleles^3^. We show how our work can be placed in the context of transmitted and untransmitted alleles in the Supplementary Information.

### Multi-locus simulations

We conducted a simple forward-in-time simulation study using realistic LD to confirm our theoretical calculations for multiple loci. We create genotypes and phenotypes of 32,000 unrelated individuals based on about 50,000 single nucleotide polymorphism (SNPs) randomly selected from the 1000 genomes project. These individuals are the parents for the next generation, creating two offspring in each of the 10 generations. We create mate assortment by ordering the phenotypes according to a specified phenotypic correlation, *ρ*_*Y*_, which is a simplified and not realistic process. However, our aim was to simply confirm our theory and to demonstrate the directions of the (co)variances and the degree of confounding under AM. More details on the simulation are given in the Supporting Information.

The left top panel of Figure 2 shows that the expected theoretical variance matches the estimated variance spread over 1000 causal loci for a wide range of variance scenarios:(i) only direct effects (V1), (ii) direct and imprinting effects (V2), (iii) direct, maternal and paternal effects (V3), (iv) direct, maternal, paternal and imprinting effects (V4) and (v) direct, maternal, paternal and imprinting effects including correlations between them (V5) under random mating. As already demonstrated numerically in Equation 2, the population level variance increases when indirect genetic and imprinting effects are present. If there are no parental influences apart from parent-of-origin effects and no sibling effects, then the variance simply increases by 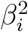, with no covariance between imprinting and direct effects. If parental genetic effects are also present, covariances then emerge and contribute substantially to the population-level variance. These covariances will be dependent upon the direction and magnitude of the maternal, paternal and imprinting effects of a given locus. The results given in Figure 2 clearly show that components of variance cannot be simply separated into independent direct and indirect sources that then simply sum. This highlights again the confounded nature of direct, indirect genetic and parent-of-origin effects.

**Figure 2:**
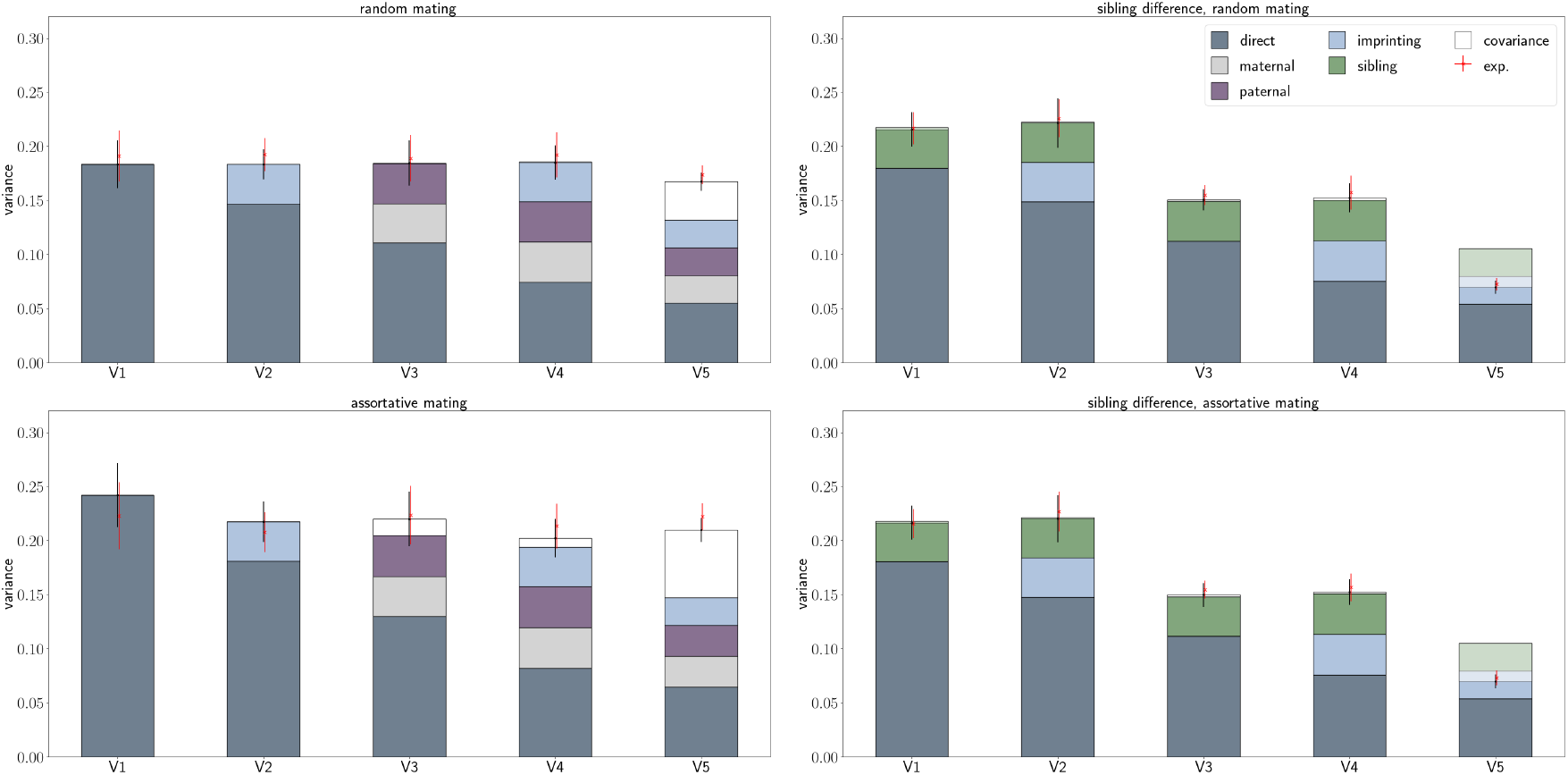
Results of simulation study. Expected theoretical total variance (exp.), and estimated variance (bars) for children (left) and sibling differences differences (right) under random mating (top) and assortative mating (*ρ*_*Y*_ = 0.25, bottom) are shown for various variance-covariance scenarios (V1-V5) for multiple loci. Bars and points represent the mean across 10 simulations, while the uncertainties indicate 2x standard deviations. Note that the direct-sibling covariance is negative and is thus shown opaque, covering parts of the variance.

Also under assortative mating for a correlation between the parental phenotypes of *ρ*_*Y*_ = 0.25, the estimated and expected theoretical variances match for the different variance scenarios, as shown in the left bottom panel of Figure 2. If we consider loci with only direct effects (V1), the variance is enlarged only due to the increased homozygosity of causal loci that is expected under AM.

As soon as indirect parental effects exist, the correlation between parental phenotypes generates a genotypic correlation between the parents which in turn creates a covariance term between the maternal and paternal effect genotypic values, *X*_*m*_*β*_*m*_*X*_*f*_ *β*_*f*_, as can be clearly seen in V3 comparing the top and bottom left panels of Figure 2. This term is an important component of variance and can be interpreted as a measure of the similarity of the indirect genetic effects among parents on their children. This suggests that assortative mating on a trait may also create assortment for parental characteristics, where biological parents create similar environments for their children, creating covariance in the indirect genetic effects that may have a large role to play in shaping the population-level variation. Until now this covariance has been generally overlooked. We propose that shared parental effects across traits can also create the cross-trait assortative mating patterns that are observed in the human population^21^.

The right panels of Figure 2 shows the expected theoretical and estimated variance when modelling the difference between siblings and including indirect sibling effects under random (top) and assortative mating (bottom). As demonstrated numerically in Equation 3, parental effects do not contribute to the variance as they cancel, although we note that with parental genetic effects the correlation within and among loci under AM alters, which changes the variance as compared to the case when parental genetic effects are absent. However, sibling effects and their covariance with the direct effects do contribute and are indistinguishable from the direct effects here, as we are calculating the difference between siblings. Interestingly, the decrease in variance under AM in case of the sibling difference is more substantial than the increase of variance in trios, as is already expected when comparing Equations 4 and 5.

## Discussion

Here, we provide a quantitative genetics model that includes the influence of indirect maternal and paternal, and parent-of-origin genetic effects on the additive genetic variance. We explicitly derive the influence of correlation of direct genetic, indirect genetic and epigenetic effects on how AM shapes the population-level genetic variance. Previous theory has established that an equilibrium is reached under AM, where there is inflated genetic variance, a larger correlation between relatives for traits driving AM and increased homozygosity at causal loci^19,22^. But imprinting and indirect genetic effects, alongside their covariances, and the change in mating frequency, have not been considered, when calculating the expected change in genetic variance.

When modelling sibling differences instead of child-mother-father trios, we additionally considered indirect genetic effects from siblings. This design removes indirect parental effects from the variance, but alters the variance of direct genetic effects in the presence of indirect sibling genetic effects. This design seems to be able to separate parent-of-origin effects under random mating. Under assortative mating, the variances in the trio study design increase, but the variances in the sibling design decrease. Therefore, we suggest that contrasting the estimates across within-family studies may give the upper and lower bounds of the degree to which the components influence the phenotypic variance within the human population.

Our results demonstrate that it is important to control for indirect parental effects when estimating the genetic variance. But even single-locus, marginal estimates common to genome-wide association studies (GWAS) from within-family studies are difficult to interpret causally^12,13^ due to the highly correlated nature of markers and effects. Controlling for the covariances among loci across the genome (both cis- and trans-correlations) when estimating genetic effects would require fitting all variants and all forms of genetic effect (direct, maternal, paternal, parent-of-origin) jointly. We show how covariances depend upon the direction and magnitude of the direct, maternal, paternal and imprinting effects within and across loci. These covariances contribute substantially to the population-level phenotypic variance, if parental genetic effects are present. In particular, covariance of the parental indirect genetic effects, a measure of the similarity of the indirect genetic effects among parents on their children, may be an important component of variance. We propose that assortative mating on a trait may also create assortment for parental characteristics, where biological parents create similar environments for their children, creating shared parental effects across traits and cross-trait assortative mating^21^.

The separation of imprinting and indirect genetic effects becomes more complex when one considers the underlying potential causal mechanism. Genomic imprinting has been considered as a form of parental effect in the sense that there is parental influence on the genotype-phenotype relationship in the offspring; while others suggest that because the offspring genotype accounts for all of the genetic variance in the offspring trait, imprinting is a form of direct effect as there is no variation explained by the parental genotype. Differentiating the genes in the parental genome that cause imprinting from parental genetic effects and from the genes in the offspring genome that become imprinted is difficult - if not impossible - especially if there are parental loci that control the imprinting state of other loci in the offspring. This complex phenomenon creates variation in offspring traits that is dependent on the combination of parental and offspring genotypes and it is important to recognize that: (i) patterns interpreted as indirect genetic effects can result from genomic imprinting (and vice versa), and (ii) it is not straightforward to distinguish parental effects on imprinting from imprinted genes showing parental expression; and (iii) under assortative mating, which is common for many human phenotypes, obtaining accurate estimates will be an even more difficult task.

We have used the terms parent-of-origin and the epigenetic phenomenon of imprinting interchangeably, assuming that parent-of-origin effects are underlain by epigenetic mechanisms, where a parent influences the genotype-phenotype relationship of their children. However, we appreciate that other factors may create parent-of-origin effects in the population^23^. Additionally, the *ρ* term used throughout this work reflects a correlation among mates at causal SNP variants (or those correlated with an underlying causal variant). A non-zero *ρ* term can be induced through actively choosing mates similar (or dissimilar) to oneself at specific phenotypic characters, termed direct assortative (disassortative) mating, which creates a phenotypic correlation among parents that translates to a correlation in underlying causal variants. Correlations can also appear through cultural, environmental, or geographical stratification in mating patterns that correlate with trait and trait-associated allele frequency stratification in the population. We do not discriminate between these here as they are likely inseparable in population-level data. Rather, we focus on demonstrating how correlations between mates at causal loci influence the phenotypic variance attributable to the DNA.

Here, we ignore dominance, genotype-environment interaction, genotype-environment correlation, epistasis, etc. Future work can incorporate these into our model. Moreover, the framework used here could be combined with mating frequencies expected under different forms of mate choice and thus be used to explore how indirect and epigenetic effects shape phenotypic variation in other systems.

In summary, population-level variance in human phenotypes is shaped by potentially complex relationships between direct, indirect genetic and imprinting effects that render their separation and quantification almost impossible, especially when assortative mating occurs within the population. Statistical modelling developments are required if we are to understand the genetic basis of human complex traits in the presence of indirect parental and imprinting effects.

## Acknowledgements

We thank members of the Medical Genomics group at ISTA for their comments, which improved this manuscript. This work was funded by an SNSF Eccellenza Grant to MRR (PCEGP3-181181), and by core funding from the Institute of Science and Technology Austria.

## Author contributions

IK and MRR conceived and designed the study, derived the theory, conducted the analyses and wrote the paper.

## Author competing interests

MRR receives research funding from Boehringer Ingelheim for work unrelated to that presented here. IK declares no competing interests.

## Code availability

The code used to simulate data can be found at https://github.com/medical-genomicsgroup/familyMC

## Supplementary Material

### Single-locus model under random mating

We first consider a single autosomal locus with two alleles (*A*_1_ and *A*_2_) with frequencies *q*_1_ and *q*_2_ that can have the following four effects on the child’s phenotype: direct (*β*_*c*_), indirect maternal (*β*_*m*_), indirect paternal (*β*_*f*_) and parent-of-origin (imprinting, *β*_*i*_). The alleles at the locus are ordered, referring first to the maternally inherited allele and second to the paternally inherited one, i.e. *A*_*m*_*A*_*f*_. The genotypes, *X*_*c*_, *X*_*m*_ and *X*_*f*_, are coded as 0, 1 and 2 for *A*_1_*A*_1_ homozygotes, *A*_1_*A*_2_ or *A*_2_*A*_1_ heterozygotes and *A*_2_*A*_2_ homozygotes, respectively. For heterozygous children, the parent-of-origin effect is positive for maternally inherited *A*_2_ and negative for paternally inherited *A*_2_. The possible genotypic and phenotypic values of the child, *Y*_*c*_, given the genotype of the parents and imprinting are shown in Table 1, where the phenotypic values represent the deviations from the case where both parents and child have *A*_1_*A*_1_ genotypes.

We begin by assuming random mating and Hardy-Weinberg equilibrium, where (*q*_1_ + *q*_2_)= 1 = (*q*_1_ + *q*_2_)^2^. The mean phenotypic value over the population is calculated as

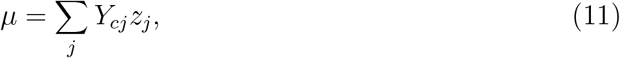

where j loops over all possible genotypes of the child, *Y*_*c*_ is the expected phenotype produced by a given genotype and *z* represents the corresponding mating frequency. The means for the different genotype classes of the child and the total mean are determined to be

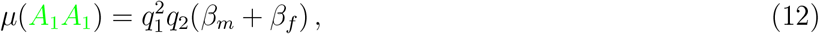

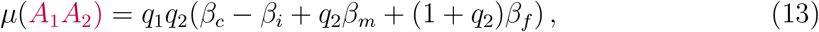

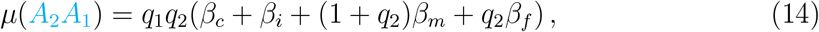

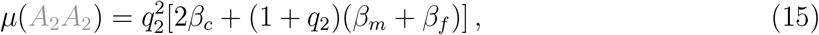

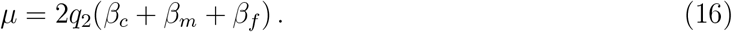

The variance is calculated as

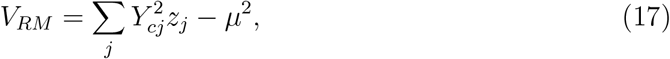

where

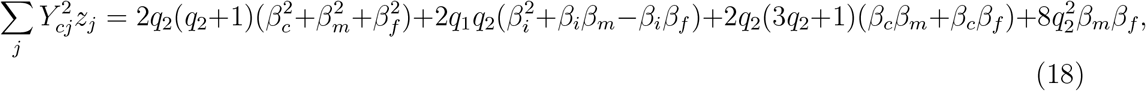

and results in

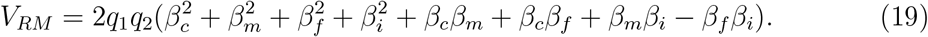

### Including indirect sibling effects

We first extend our single autosomal locus model to the case of siblings where there are the following five effects on the child’s phenotype: direct (*β*_*c*_), indirect maternal (*β*_*m*_), indirect paternal (*β*_*f*_), parent-of-origin (imprinting, *β*_*i*_) and sibling (*β*_*s*_). The possible genotypic and phenotypic values of the child, *Y*_*c*_, given the genotype of the parents, sibling and imprinting are shown in Table 2, where the phenotypic values represent the deviations from the case where all parents and children have *A*_1_*A*_1_ genotypes.

We begin by assuming random mating and Hardy-Weinberg equilibrium, where the means for the different genotype classes of the child and the total mean are determined to be

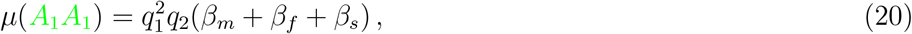

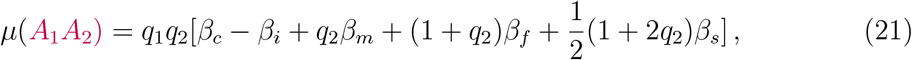

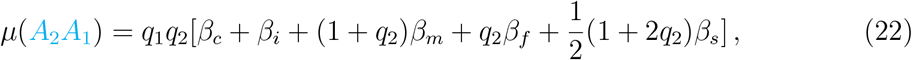

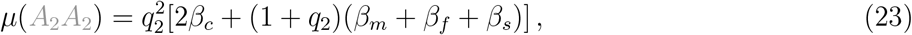

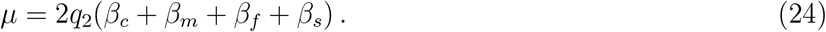

The variance is calculated as

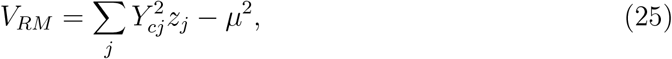

where

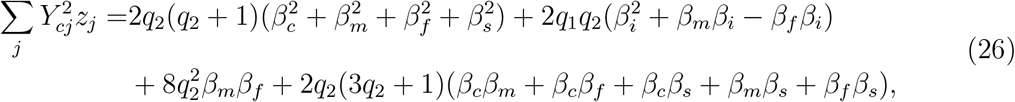

and results in

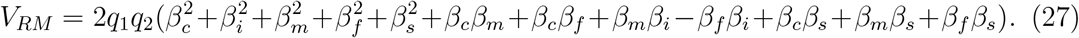

The mean and population-level variance depend on the mating frequency, *z*, of the parents. We again introduce a correlation *ρ* between the maternal and paternal alleles which impacts the mating frequencies, *z*_*AM*_, as can be seen in Table 1. The means for the different genotype classes of the child and the total mean are now determined to be

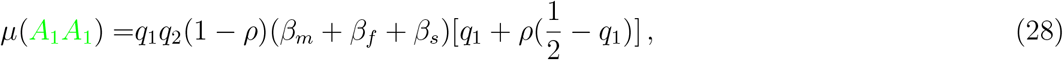

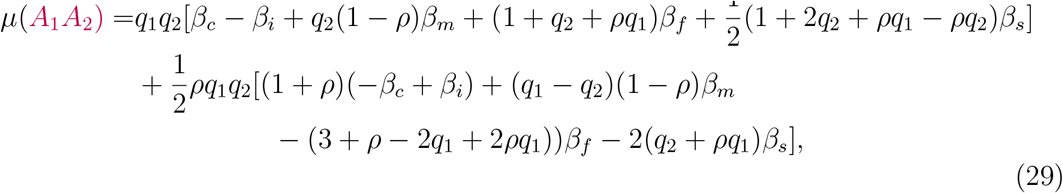

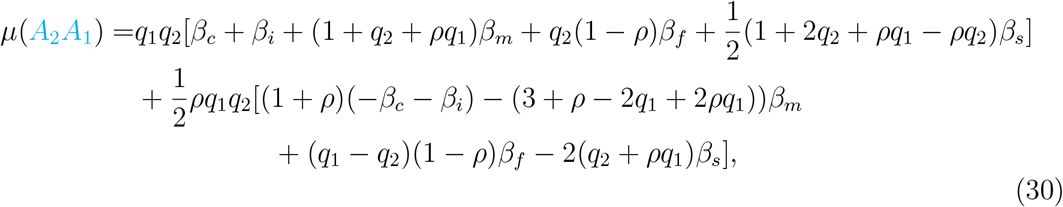

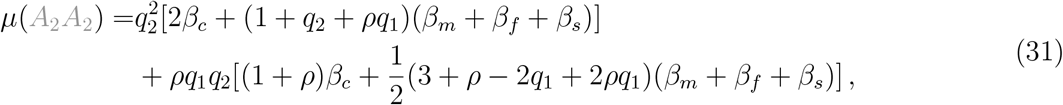

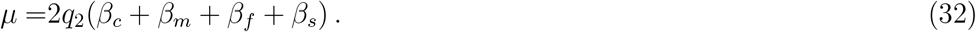

The genotypic mean is the same as the one assuming that there are no correlations between the parental genotypes, which is expected as the allele frequencies do not change. The variance is then

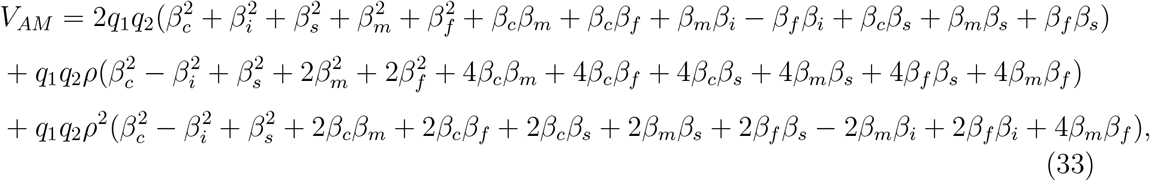

using

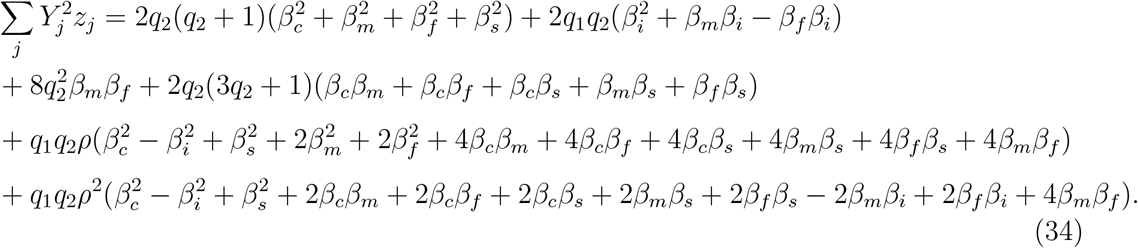

This shows that indirect effects of siblings are also confounded with the direct effects, the parental genetic effects and the imprinting effects. Assortative mating acts to inflate the covariance of the indirect sibling genetic effects and all other components, making a partitioning of variance components intractable.

## Modelling the phenotypic difference of siblings

Previous studies^16^ have proposed to control for indirect effects by looking at the phenotypic and genetic difference in siblings. Table 2 shows the possible genotypic and phenotypic values of the child, *Y*_*c*_, and sibling, *Y*_*s*_, given the genotype of the parents, the respective sibling and imprinting, as well as the phenotypic difference between the offspring, *Y*_*c*_ *− Y*_*s*_.

The means in phenotypic difference for the various genotype classes of the child and the total mean are determined to be

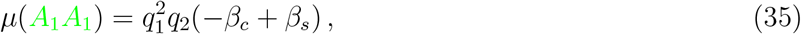

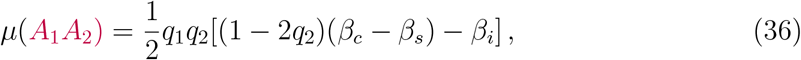

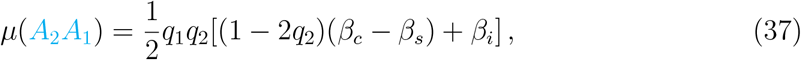

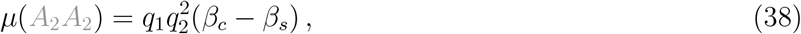

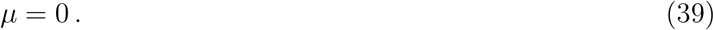

The total mean is 0 as expected when calculating the difference between siblings. Thus, the variance is

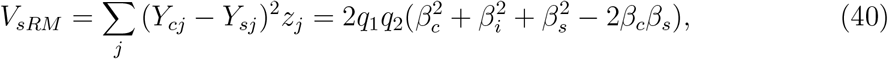

which shows that indirect parental effects do indeed cancel in the difference, but indirect sibling and imprinting effects remain.

### Single-locus model under assortative mating

The expected phenotype of an offspring of a given genotype depends on the probability that it was produced by each of the different possible parental genotypes. Assortative mating (AM) acts to alter the mating frequencies in the population as it increases parental similarity at trait-associated loci which is reflected in the correlation, *ρ*, among the parents in their alleles. This changes the genotype frequencies, but does not affect the minor allele frequencies in the population. The genotype frequencies under AM are given in Ref.^15^. The means for the different genotype classes of the child and the total mean are determined to be

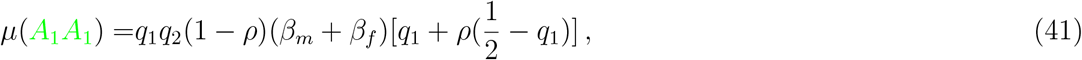

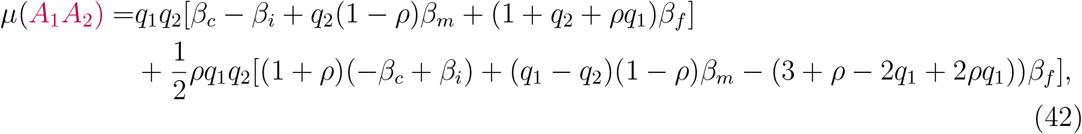

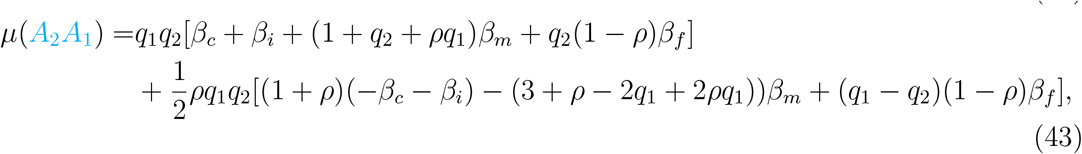

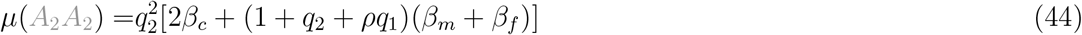

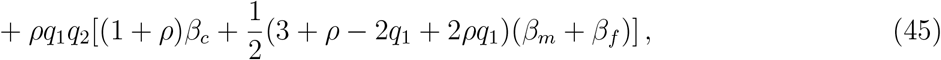

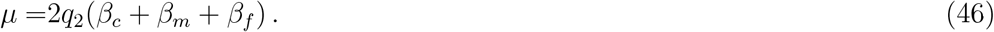

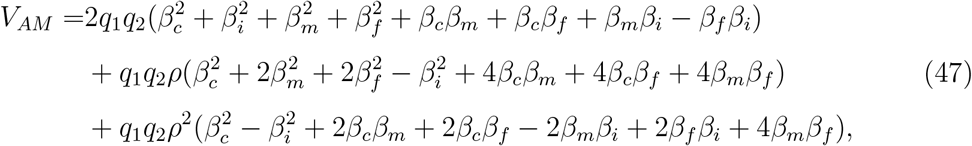

using

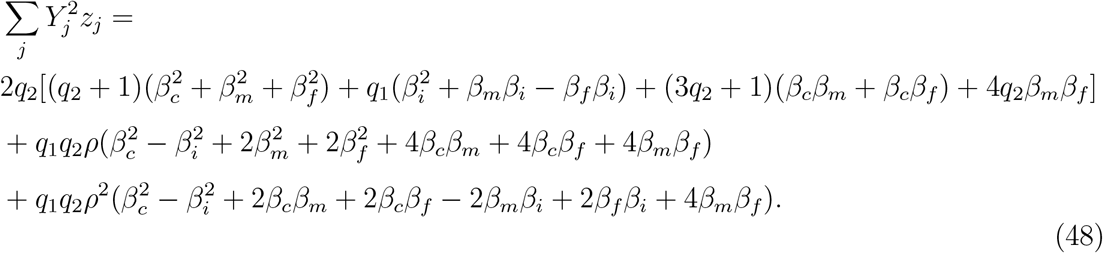

All terms are increased under assortative mating with the exception of imprinting which is reduced. Note that there is now an additional correlation term between the parental genetic effects, *β*_*m*_*β*_*f*_, that depends on *ρ*.

### Modelling the phenotypic difference in siblings

Extending the model to also include the change in genotype frequencies due to assortative mating, we find

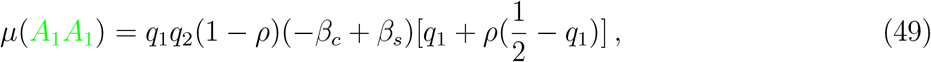

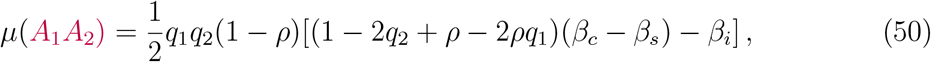

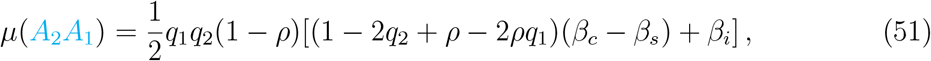

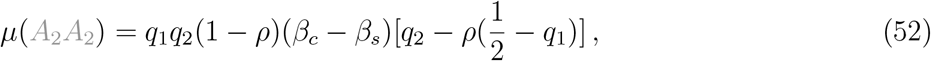

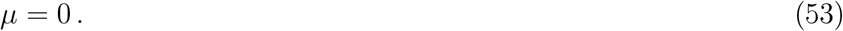

The variance is reduced by the parental correlation, *ρ*,

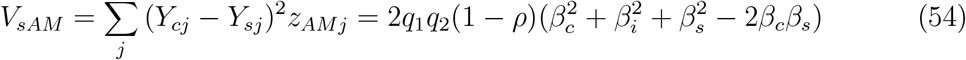

### Transmitted and untransmitted parental alleles

We sought to place our work in the wider context of previous research, which has focussed on transmitted and untransmitted alleles ^3^. The phenotype of a child, *Y*_*c*_, that is influenced by direct, indirect maternal, indirect paternal and imprinting effects across *l* loci, can be written in terms of transmitted (*A*_*m*_, *A*_*f*_) and untransmitted maternal and paternal alleles (*A*_*\m*_, *A*_*\f*_):

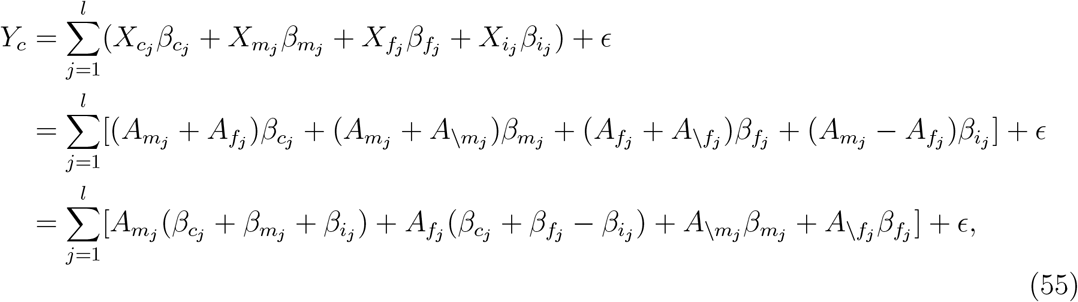

where *ϵ* represents the residual error due to other environmental influences. Note that the imprinting effect will have a positive sign if only *A*_*m*_ is transmitted, a negative one if only *A*_*f*_ is transmitted, and will be equal to 0 for all the other cases. Thus, a transmitted allele has direct, indirect and imprinting effects, while an untransmitted allele only has indirect effects on a child’s phenotype.

Assuming Hardy-Weinberg equilibrium and random mating, thus that the four alleles are independent, the total variance that each transmitted and untransmitted allele contributes to the trait of a child per locus *j* is

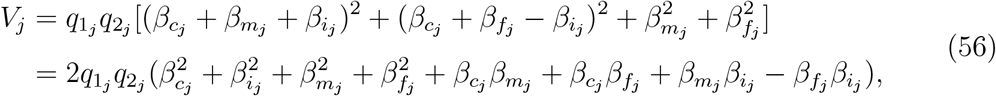

where 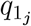 and 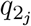are the allele frequencies. Equation 56 is equivalent to Equation 2.

When effect sizes are estimated in GWAS, a linear relationship between an individual’s phenotype and their genotype, i.e. the two transmitted alleles, is assumed, ignoring imprinting effects. Thus, the SNP heritability, defined as the proportion of phenotypic variance among children attributable to the SNP genotypes of the children, would correspond to

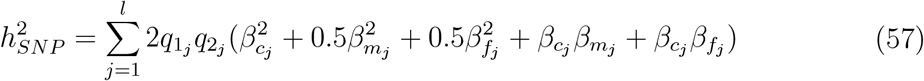

assuming that all loci are independent. Our results suggest that in reality, it is not known how much the indirect effects contribute to the GWAS estimate unless all the direct and indirect terms are taken into account, as we do not know the size and magnitude of the covariances between direct and paternal effects, nor do we know if indirect components are actually totally estimated or estimated as part of the residual variance, nor is it clear how the abundant covariances across loci will influence the effect estimates (both cis- and trans-LD). It is also important to note that the variance due to the untransmitted alleles does not equal the total variance of the indirect effects, but only half (at most). Thus, the effects attributed to the untransmitted alleles are in fact not a good estimate of indirect parental effects, as has already been noted by Ref. ^13^.

Previous work^3^ used polygenic risk scores computed from transmitted alleles, 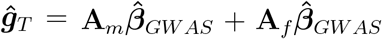, and untransmitted ones 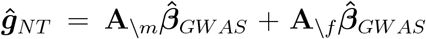, to separate direct from indirect effects. The effects of transmitted and untransmitted alleles,*θ*_*T*_ and *θ*_*NT*_, are estimated from the linear regression,

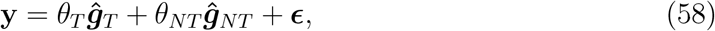

where **y** is a vector of phenotypic observations and **y**, ĝ_*T*_ and ĝ_*NT*_ are all standardized to have mean zero and variance 1. The direct effects of the transmitted polygenic risk score, δ, are then determined by calculating the difference between transmitted and untransmitted effects, δ = *θ*_*T*_ *− θ*_*NT*_. The estimated variance accounted for by the direct effect alone is then given by

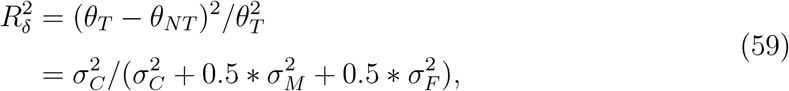

where 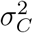 is the variance contributed by the direct effects 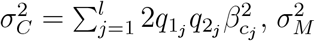 is the variance contributed by the indirect maternal effects 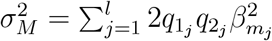, and 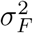 is the variance contributed by the direct effects 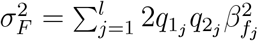

Equation 59 makes the following assumptions: 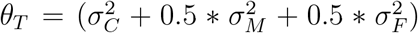 and 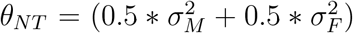 (ii) loci are independent; (iii) there are no covariances among direct and indirect effects; (iv) imprinting effects are absent; (v) transmitted and untransmitted alleles have the same indirect effects; and (vi) estimation error can be ignored. Note that the estimated variance for direct effects, 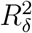, in Equation 59 is defined differently than the direct variance, 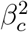, in Equations 2-5. 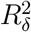 is a relative estimate of the direct effects in the transmitted predictor, while 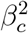 is an absolute value and can only be transformed to something similar to 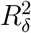 by making an assumption on the heritability of the transmitted alleles.

### Forward-in-time simulation

A simple, but realistic simulation (based loosely on Ref. ^24^) for parents and children is set up and moved forward in time for a number of generations, *ngen* = 10, to simulate assortative mating. Genotypes of 32,000 unrelated individuals in generation 0 are created with a realistic LD based on 52,310 SNPs. The SNPs were randomly selected across all chromosomes from the 1000 genomes project (http://ftp.1000genomes.ebi.ac.uk/vol1/ftp/release/20130502/) using vcfrandomsample (https://github.com/vcflib/vcflib#vcflib) to downsample the data to make it more manageable. A pair of individuals produce two offspring each by recombining the parents’ haplotypes based on a genetic map (release b37, for example from https://github.com/odelaneau/shapeit5/tree/main/resources/maps/b37). The offspring of generation 0 become the parents for the next generation. Parents after generation 0 mate based on their ordered phenotpyes to create assortative mating. The phenotypes for ordering are calculated as:

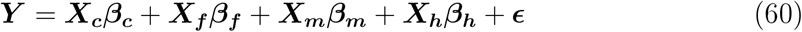

where *ϵ* is a value drawn from a Normal distribution with mean 0 and standard deviation of 0.01 to facilitate ordering. The phenotypes are ordered to introduce a correlation, *ρ* = 0.25, between the parents. This procedure is a simplified version of the unknown realistic process. For the implementation of the AM procedure, see Code availability.

Simulation studies were carried out 10 times each for the following five different variance-covariance matrices of effects,

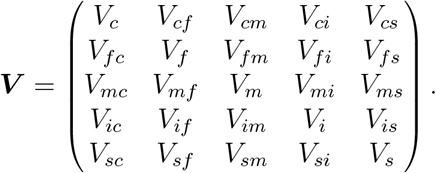

When the indirect effects of siblings are not modelled, the last column and the last row are removed.

1. Scenario 1 represents a covariance matrix with only direct effects and, in case of the sibling difference, sibling effects:

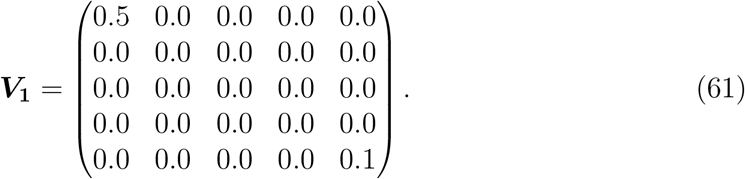
2. In the second scenario, direct as well as parent-of-origin effects and, in case of the sibling difference, sibling effects are contributing to the variance:

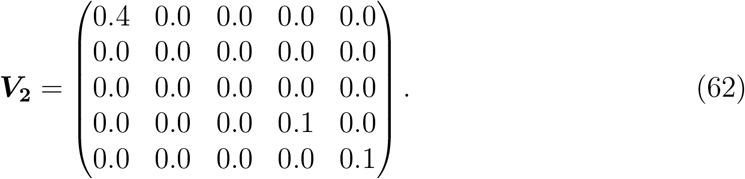
3. Scenario 3 assumes direct effects as well as smaller indirect maternal and paternal effects and, in case of the sibling difference, sibling effects:

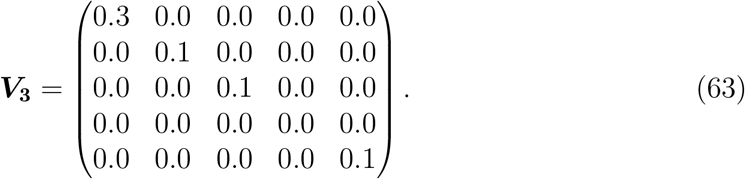
4. In scenario 4, the total variance of 0.5 is spread over all four genetic components. In the case of the sibling difference, there are additionally sibling effects. There are no correlations.

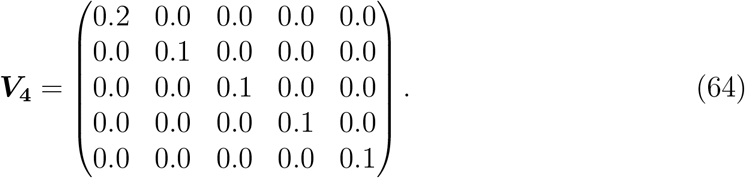
5. The last scenario represents a scenario where all effects are correlated:

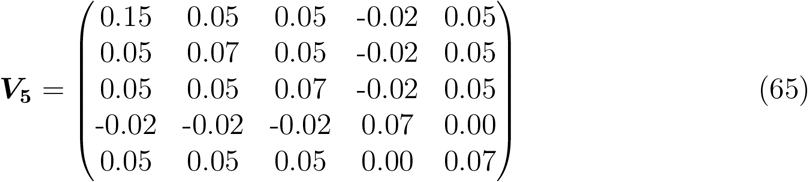

Effects were generated for 1000 markers with a multivariate normal distribution with mean 0 and variance **V** scaled by the number of causal markers and randomly assigned to 1000 markers for each simulation. The effects were assumed to be constant across generations.

## Notes

### Summary of Updates

We updated our simulation work that accompanies the models to include more loci to better reflect a polygenic trait architecture. We clarify that by confounded, what we meant was that estimating and seperating indirect genetic effects and parent-of-origin effects is difficult for the reasons explain within the paper.

